# Cueing memory during sleep is optimal during slow-oscillatory up-states

**DOI:** 10.1101/185264

**Authors:** Maurice Göldi, Eva van Poppel, Björn Rasch, Thomas Schreiner

**Author notes:** Maurice Göldi, University of Fribourg, Department of Psychology, Division of Cognitive Biopsychology and Methods, Rue P.-A.-Faucigny 2, CH-1701 Fribourg, Switzerland. Eva van Poppel, University of Fribourg, Department of Psychology, Division of Cognitive Biopsychology and Methods, Rue P.-A.-Faucigny 2, CH-1701 Fribourg, Switzerland. Corresponding authors: Thomas Schreiner, Radboud University, Donders Institute for Brain, Cognition and Behaviour, Kapittelweg 29, 6525 EN, Nijmegen, The Netherlands, Björn Rasch, University of Fribourg, Department of Psychology, Division of Cognitive Biopsychology and Methods, Rue P.-A.-Faucigny 2, CH-1701 Fribourg, Switzerland.

## Abstract

Slow oscillations play a major role in neural plasticity. It is assumed that slow oscillatory up-states represent crucial time windows for memory reactivation and consolidation during sleep. Here we experimentally tested this assumption by utilizing closed-loop targeted memory reactivation (closed-loop TMR): Healthy participants were re-exposed to prior learned foreign vocabulary during up- and down-states of slow oscillations, respectively, in a within-subject design. We show that presenting memory cues during slow oscillatory up-states robustly improves recall performance, whereas memory cueing during down-states did not result in a clear behavioral benefit. On a neural basis successful memory reactivation during up-states was associated with a characteristic power increase in the theta and sleep spindle band. Such increases were completely absent for down-state memory cues. Our findings provide experimental support for the assumption that slow oscillatory up-states represent privileged time windows for memory reactivation, while the interplay of slow oscillations, theta and sleep spindle activity promote successful memory consolidation during sleep.

---

The consolidation of memories critically depends on hierarchically nested oscillatory brain mechanisms. It has been proposed that the systematic interaction of neocortical slow oscillations (SOs), thalamic sleep spindles and hippocampal sharp wave ripples (SWRs) during sleep reflect the mechanistic vehicle of memory reactivation and thus consolidation during sleep ^1^. The <1 Hz SO represents the most prominent signature of slow wave sleep (SWS) and is generated in neocortical and thalamic circuits. SOs comprise alternations between periods of neuronal membrane hyperpolarization, accompanied by widespread neuronal silence (“down-states”), followed by depolarized neuronal “up-states”, accompanied by sustained firing ^2^. Critically, SOs are thought to coordinate spontaneous memory reactivation processes during sleep, by providing the temporal frame for active memory consolidation ^3^. It is assumed that the SO up-states drive memory reactivation in the hippocampus together with sharp wave ripples and thalamo-cortical sleep spindles ^4^. The formation of spindle-ripple events is suggested to enable the hippocampal-neocortical dialog and the redistribution of reactivated hippocampal memory information to neocortical long-term stores^5,6^.

Despite these theoretical predictions, the mechanistic role of SO up-states for memory reactivation during sleep in humans remains ambiguous. In general, the functional significance of sleep-related memory reactivation in humans has been demonstrated by a series of studies showing that inducing reactivation processes experimentally (targeted memory reactivation; TMR) improves the consolidation process and thereby affects subsequent recall performance ^7^. TMR studies follow the rationale that memory cues associated with prior learning are presented again during subsequent non rapid eye movement (NREM) sleep to trigger reactivation processes and consequently boost later memory performance. This approach has repeatedly proven successful for context cues such as odors ^8–10^ and for specific item cues such as sounds ^11–13^, melodies ^14–16^ or verbal material ^17–19^. Importantly, all of these studies presented the memory cues at random points in time during NREM sleep, without taking the on-going oscillatory activity, specifically the phase of SOs, into account. Given the assumed role of SO up-states in driving memory reactivations during sleep, we hypothesized that experimentally aligning the memory cues to the initiation of SO up-states (negative-to-positive transition of the surface slow-wave; see ^20,21^) should be critical for successful TMR, resulting in improved retrieval performance after sleep. In contrast, presenting memory cues at the onset of the SO down states (positive-to-negative transition in the surface slow-wave) should block the memory benefit of TMR. We therefore used SO phase-specific targeted memory reactivation (closed-loop TMR) to test the functional role of SO up-states for memory consolidation in humans. To investigate how closed-loop TMR influences the reactivation and consolidation of memories, we used a vocabulary learning task ^17,19^. After learning 120 Dutch-German word pairs, 16 healthy young participants slept for 3 hours in the laboratory (see ‘Materials and Methods’ section, for details). We applied an online SO detection algorithm to present subsets of the prior learned words either during the presence of SO upstates or down-states. As a control condition some of the prior learned words were not replayed at all (uncued words). After sleep participants were tested on memory for the German translations by a cued recall procedure (see Fig. 2a, for a summary of the procedure). We show that words presented during SO up-states were associated with a robust improvement in recall performance compared to uncued words. On a neural basis, successful up-state TMR was associated with characteristic cue-related increases in theta and sleep spindle power. Words replayed during SO down-states did not show this distinct oscillatory pattern of successful memory reactivation and also no significant performance improvement.

**Figure 2.**
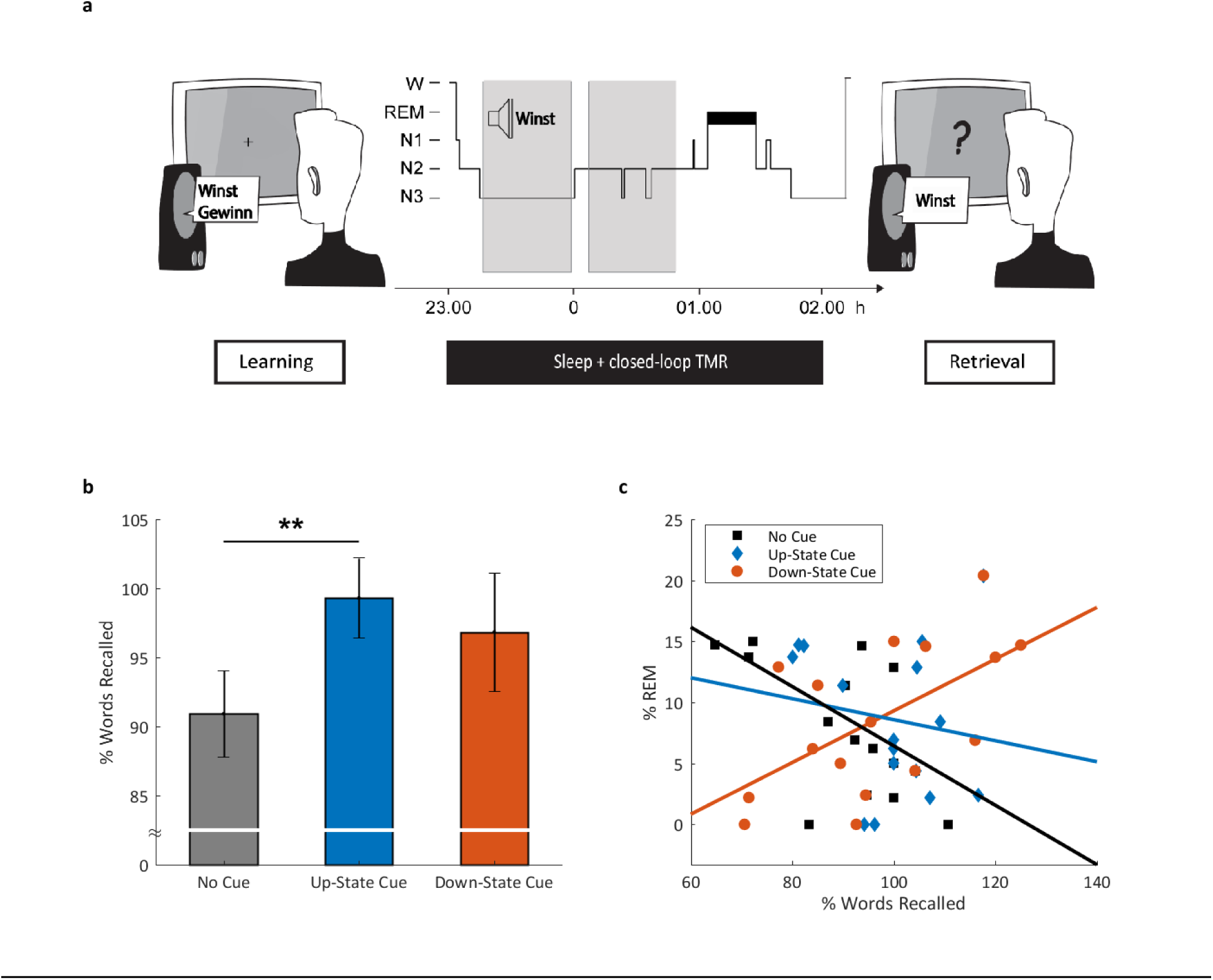
Experimental Procedure and memory task results. a) After studying 120 Dutch-German word pairs in the evening, participants slept for 3 hours. During NREM sleep, 40 Dutch words were presented during SO up-states and 40 Dutch words were presented during SO down-states using closed-loop TMR. 40 Dutch words were not replayed. A cued recall procedure was applied after sleep, testing the participant’s memory for the German translations. b) Presenting prior learned words during SO up-states significantly enhanced memory performance compared to uncued words. Recall performance of words replayed during SO down-states did not differ from the two other categories. Retrieval performance is indicated as percentage of recalled German translations with performance before sleep set to 100%. Values are mean ± s.e.m. **P ≤ 0.025. c) Correlation between memory performance and relative time spent in REM sleep. Memory performance for words presented during down-states is positively correlated with time spent in REM sleep (r(14) = 0.59, P = 0.017). There was no significant correlation for words presented during up-states (r(14) = −0.16, P = 0.552), and a marginal significant negative correlation for uncued words (r(14) = −0.49, P = 0.052).

## Results

### Accuracy of the Closed-Loop Algorithm

We first examined whether our algorithm correctly distinguished between words replayed during up-and down-states. We calculated ERPs separately for cues presented in the down-to-up phase transition of the cortical slow wave (targeted area for the up-state cues) and for cues presented in the up-to-down-phase transition (targeted area for the down-state cues, Fig. 1a). As expected, the ERP analysis confirmed that up and down state cues were played at highly distinct times of the cortical slow wave and targeted the expected areas (Fig. 1b; see Supplementary Fig. 2 for ERPs differentiated by remembered and non-remembered words). Despite the pre-stimulus peaks having clearly opposite polarities, post-stimulus ERPs of the two cueing conditions followed a similar temporal evolution (see supplementary section ‘Algorithm Accuracy’ for further details). To further assess the accuracy of the SO detection algorithm, we determined the phase of the cortical SO at stimulus onset. On a subject level, up-state cues were associated with a mean phase angle of 338.60° ± 20.46°, while down-state cues had a mean phase angle of 132.31° ± 18.46° (see Fig. 1c, top row and Fig. 1c, bottom row, for results at the trial-by-trial level). Thus, the onset of our memory cues corresponded very accurately with the early phases of the targeted areas, assuring that mostly the whole word length (~400 ms) was played during the intended SO state (see Fig. 1a for an overview and Supplementary Fig. 4 for a phase analysis of each individual subject).

**Figure 1.**
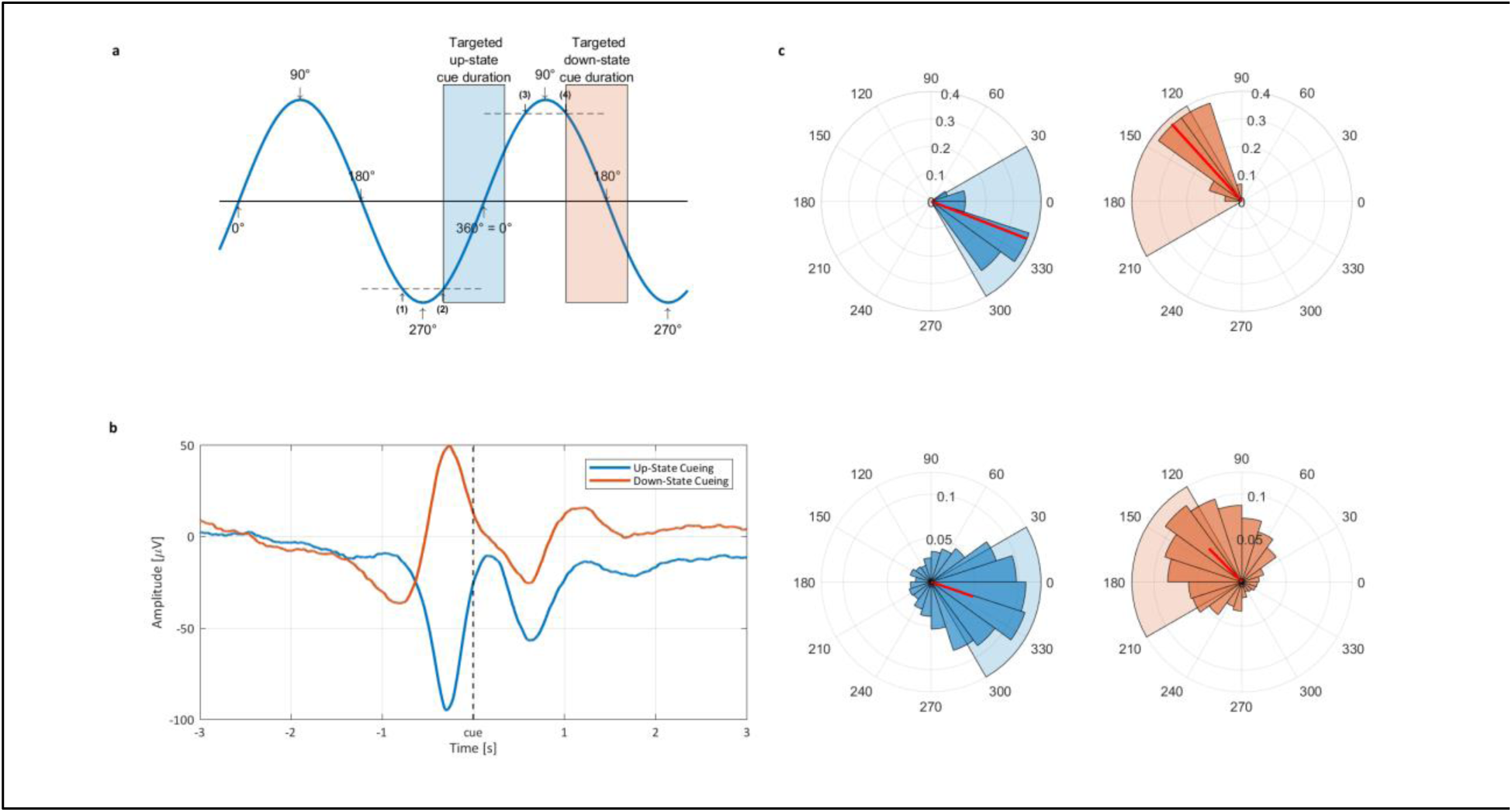
Closed-loop TMR algorithm evaluation. a) Schematic overview of the slow-wave detection algorithm and the targeted areas for up-state TMR (blue) and down-states TMR (red). b) The ERP-analyses revealed that up-state cues were located at the down-to-up transition of the cortical slow wave (beginning of slow oscillatory up-state), and that down-state cues were played at the up-to-down transition (beginning of slow oscillatory down-state). c) Phase angle at stimulus release. The top row illustrates the angles averaged at subject level, the bottom row shows results on trial level. The left column indicates the up-state phases, while the right column shows the down-state phase angles. All data is shown for electrode Fz.

### Behavioral Results

As predicted, we observed a robust improvement of memory performance when word-cues were presented during the up-state of the slow oscillation (see Fig. 2b): After the sleep interval participants remembered 99.3 ± 2.89% of the prior learned words which were presented during SO up-states, whereas they remembered only 90.92 ± 3.14% of the uncued words (t_15_ = 2.62; *P* = 0.019, two-tailed). In contrast, memory for words presented during SO down-states (96.83 ± 4.27%) did not differ from uncued words (t_15_ = 0.93, *P* = 0.366). On a descriptive level, memory performance for words presented during down-states was just in-between up-state and uncued words. Furthermore no difference to memory performance associated with up-state cued words was observable (t_15_ = 0.43, *P* = 0.673). It must be noted that the variance was also descriptively higher for words cued during the down-state than the other two categories (down: 292.29; up: 133.20; non-react: 157.58). In addition, down-state cued words were negatively correlated with memory for uncued words (r(14) = −0.45, *P* = 0.079). This coefficient differed significantly (Z = 2.34, *P* = 0.015) from the positive correlation between up-state cued words and uncued words (r(14) = 0.44, *P* = 0.091). Consequently, the overall multivariate analysis of variance including all three word categories simultaneously only reached a statistical trend (F_(2,14)_ = 3.21, *P* = 0.071). Furthermore, we explored the associations between memory performance and time spent in the different sleep stages (for descriptive values of sleep stages, see Table 1). Interestingly, only participants with high amounts of REM sleep profited from down-state cueing, while participants with low or no REM sleep did not (r(14) = 0.59, *P*= 0.017; see Fig. 2c; please note that no cue was presented during REM sleep). We observed no other significant correlations between any sleep stage and memory performance in any word category (all *P* > 0.05). We also observed no significant correlation between the number of cues played during NREM sleep and memory performance, neither in the up-nor down-state category (both, *P* > 0.180). Descriptively, each person received 308.94 ± 18.98 cues during the night, which corresponds to about 5 repetitions of all cued words, with 50.29 ± 0.21% up-state cues and 49.71 ± 0.21% of the words presented during down states.

**Table 1.**
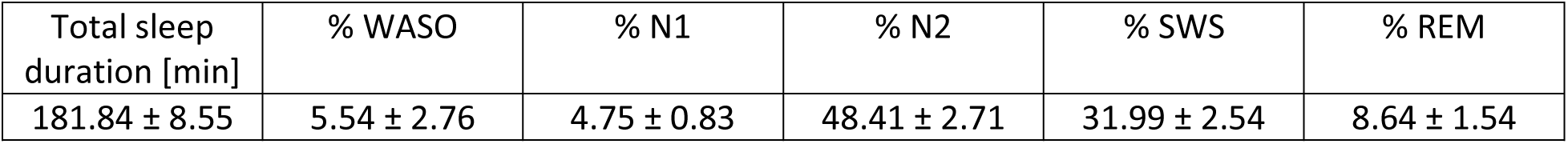
Sleep parameter. N1, N2: NREM sleep stages N1 & N2, SWS: slow-wave sleep (N3), REM: rapid eye movement sleep, WASO: wake after sleep onset. Values are means ± s.e.m.

| Total sleep duration [min] | % WASO | % N1 | % N2 | % SWS | % REM |
| --- | --- | --- | --- | --- | --- |
| $181.84 \pm 8.55$ | $5.54 \pm 2.76$ | $4.75 \pm 0.83$ | $48.41 \pm 2.71$ | $31.99 \pm 2.54$ | $8.64 \pm 1.54$ |

### Oscillatory Results

Based on our previous reports ^17,19^, we focused the time-frequency analysis on oscillatory power in the theta band (5-8 Hz) and the sleep spindle band (11-15 Hz) between the time points 0 ms and 2000 ms after stimulus onset. For memory cues played during the up-state we observed a significant increase in theta power for later remembered compared to later non-remembered words between 920 and 1480 ms involving a cluster of 22 channels (*P* = 0.05, see Fig. 3a). The significant electrodes had a right central distribution (see Fig. 3b left column, bottom row). Also in the spindle band, the overall analysis revealed a significant increase in spindle power for remembered compared to non-remembered words between 830 and 1770 ms involving all 31 electrodes (*P* = 0.023, see Fig. 3b left column, top row). In contrast to cues presented during SO up-states, we did not observe any significant power differences for remembered vs. non-remembered words played in the SO down-state, neither for the theta (*P* > 0.30) nor the spindle band (*P* > 0.60, see Fig. 3c and d). Even a more restricted test-statistics limited to the time-range of the up-state clusters revealed no significant effect (not shown; theta: *P* = 0.213, spindle: no cluster found). The general oscillatory differences between up- and down-state cueing are shown in Supplementary Fig. 5 and discussed in the supplementary results section ‘Oscillatory Analysis Up versus Down’.

**Figure 3.**
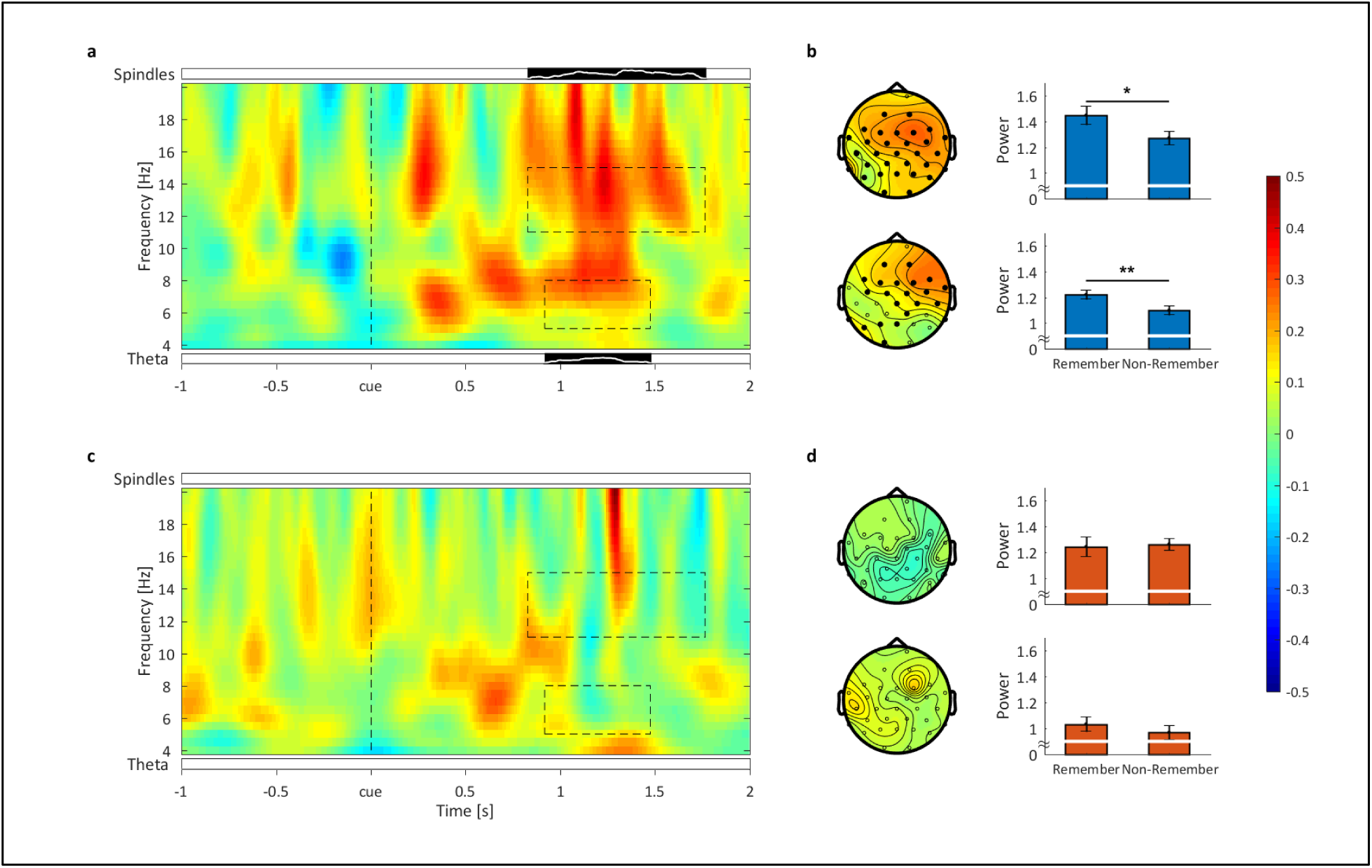
Oscillatory results. Time-frequency contrasts between remembered and not-remembered words in the theta and sleep spindle band for a) up-state and c) down-state cues, shown for the representative Fz electrode. Black bars (significant cluster in frequency band analysis) with white lines below and above the time-frequency plot show the number of significantly differing electrodes for the theta and sleep spindle band respectively. The full height of the bar corresponds to 100% (31) electrodes. Dashed boxes indicate the areas of significant difference between remembered and not remembered words. These time-windows were used to illustrate the topographical distributions (b and d) left column, top row spindle band, bottom row theta band; significant electrodes shown as filled black dots). b) and d) right column show the mean power within the significant clusters, averaged over the significant electrodes, all frequencies and time in the sleep spindle (top) and theta (bottom) band. For up-state cueing a) remembered words show enhanced power in the theta (5-8 Hz) as well as the sleep spindle (11-15 Hz) range compared to not-remembered words. Averaged over time duration, channels and frequency band, within these clusters this difference was highly significant in the theta band (t_15_ = 2.63, P = 0.01; see b right column, bottom row) and in the spindle band (t_15_ = 2.37, P = 0.016; see b, right column, top row). For words presented during down-states c) no significant difference emerged between remembered and forgotten words, neither in the sleep spindle nor the theta band. Consequently, averaged activity in those clusters observed in the analysis of SO up-states did not reveal any significant differences for down-state cues, neither in the theta (t_15_ = 1.02, P = 0.161) nor the spindle band ( t_15_ = −0.33, P =0.627). Mean ± SEM are indicated. **: P < 0.01; *: P< 0.05.

## Discussion

The present study investigated for the first time the impact of SO phase-dependent memory reactivation during sleep on memory consolidation. We used a simple, yet effective algorithm to target the presentation of prior learned words specifically into the transition between neuronal quiescence and synchronized neuronal activity and vice versa (i.e. SO up and down-states). We demonstrate that presenting memory cues during the presence of SO up-states significantly improved memory performance compared to uncued words. In contrast, cueing words during SO down-states did not exhibit such a clear beneficial effect. However, participants who spent more time spent in REM sleep after NREM cueing showed a stronger memory benefit from cues presented during down-states. Our oscillatory analyses demonstrate that successful TMR during up-states was related with higher theta and spindle band activity than non-successful TMR as described previously ^12,17–19,22^. This characteristic oscillatory pattern of successful memory reactivation during NREM sleep was not observable for the down-state condition. In our study, we targeted the SO transition from down- to up-state (down state condition) and up- to down-state (up-state condition) to specifically reactivate memories during sleep. We aimed for these transition periods as it has been shown that neuronal down-states occur slightly before the negative peak of the surface slow wave, while neuronal up-states start with the negative-to-positive transition of the cortical slow-wave ^20,21^. Our algorithm accurately presented the memory cues in the intended target areas. In addition, our ERP analysis shows that, while the signals of the two cue targets differed strongly before cue onset, they followed a similar temporal evolution after cue presentation, indicating that auditory cues pushed the brains neuronal population into similar temporal successions as measured by surface EEG, regardless of the underlying endogenous brain state.

Taken together, our results suggest that SO up-states represent an optimal time window for targeted memory reactivation and accompanying consolidation. In contrast, TMR during down-states did not ultimately block but mostly attenuated the chances of successful memory consolidation.

Several theoretical considerations ^4,5,23–26^ lead to the prediction that SO up-states drive the reactivation of memories during sleep, thereby representing a critical time window for memory consolidation. During slow oscillatory up-states, more and more neurons fire synchronously ^20^. This synchronous firing leads to a higher probability of activating a cascade of downstream–connected neurons. Memory traces are thought to be stored in the brain as interconnected neurons ^27–29^. By randomly activating a large portion of neurons during the up-state, these interconnected neurons (i.e. memory traces) fire together with a high probability ^3^. We suggest that presenting auditory cues during the presence of SO up-states, activates neurons of the associated memory trace with a heightened probability. In combination with the heightened random neural activations of the up-state, a neuronal cascade involving the complete memory trace is triggered with increased probability. Interestingly, ERP associated with up-state TMR indicated a regular, uninterrupted 1 Hz oscillation after cue onset, possibly supporting the endogenous reactivation and consolidation mechanism, while down-state TMR seemed to disrupt the ongoing SO pattern and lead to a delayed negative trough of the SO wave.

Still, presenting memory cues during the presence of a down-state might also activate the corresponding memory trace with an above chance probability. However, as the brain is at this point in a quiescent state, the reactivation of the specific memory trace might not be fully supported by the endogenous activity of the brain. The chance of reactivating a memory trace during the down-state, therefore remains inferior to up-state reactivation. Also, the natural continuation of the 1 Hz oscillation associated with down-state cueing seemed to be disturbed through stimulus presentation, leading to a slight delay in the subsequent negative ERP deflection relative to its undisturbed form. Nevertheless, both cueing time points seem to increase the chance of reactivating a memory relative to random endogenous chance reactivations. In contrast to our experimental findings, a recent analysis by Batterink and colleagues ^30^ identified the SO down-state to represent the optimal phase for TMR. In a post-hoc evaluation of two previous TMR studies ^11,31^, the authors found that the amount of forgetting was lowest for items presented just before the onset of down-states. The authors explained this rather surprising result (i.e. that the optimal phase for TMR was found to occur quite far in advance to the anticipated one, the SO up-state), by a potential time lag caused by auditory stimulus processing and concluded that a closed-loop TMR approach would shed further light on these findings. Our results indicate that the optimal phase for TMR and thereby memory reactivation is during down-to up-transition of the SO. However, general differences between the studies (e.g. utilized task, differing sound cues etc.) could potentially also account for the timing discrepancies. Our study was specifically designed to test the difference between up-state, down-state and non-cued words on memory performance. Thus, our algorithm targeted the early stage of the up-to down-and down-to up-transition and did not aim for the pre-down-state peak interval, which was found preferable by Batterink and colleagues ^30^.

An interesting finding of our current work is that cueing success for down-state TMR was positively correlated with the relative time spent in REM sleep. Two recent studies have shown, utilizing an afternoon nap design, that REM sleep systematically influenced the effect of TMR on new word learning ^32^ and the integration of newly learned spoken words ^33^. This pattern of result led the authors to assume that TMR during NREM sleep might reactivate a certain memory trace, and at the same time prepare it for integration into pre-existing associative networks during the next REM sleep cycle (which might be associated with de-stabilization of a given memory trace during NREM sleep, but see ^34^). In the light of these results we propose that memory reactivations targeted into the optimal SO up-state successfully reactivate and stabilize the memory trace immediately through the critical interplay of SO, theta and sleep spindle activity. In contrast, memory cues targeted into suboptimal (non-up-) states also have the chance to reactivate the memory trace but lack crucial stabilization processes, possibly due to the marginal spindle activity. These memory traces are therefore dependent on the re-stabilization during REM sleep. Furthermore, random TMR during an afternoon might be specifically vulnerable to presenting cues during suboptimal states due to shallower afternoon sleep than nighttime sleep.

While the role of REM sleep in stabilizing memory representations was specifically associated with down-state TMR, successful up-state TMR was directly linked with oscillatory power increases in the theta and sleep spindle range. A growing number of TMR studies ^17,18,22,35–37^ report such elevated levels of oscillatory theta and sleep spindle power to be tightly associated with cueing success, indicating a critical role of both frequency bands for the reactivation and stabilization of memories during sleep. According to our working model, increases in theta power indicate successful reactivation of the memory by cueing, whereas related increases in spindle power support the consolidation and integration of reactivated memory into cortical networks for long-term storage^38^. Here we observed significant increases in theta power for remembered as compared to non-remembered words from 0.92 to 1.48 seconds with regard to up-state cues, thus exhibiting some overlap concerning the first positive ERP-peak after stimulus onset. This temporal pattern also lies within the potentially crucial 1.5 second post-stimulus time window identified in our previous work ^17^, in which consolidation processes are prone to interference, leading to a blockade of associated reinstatement processes. There was no difference between remembered words and non-remembered words in theta activity with regard to down-state cues.

In addition, we found increase in sleep spindle power between 0.83 and 1.77 seconds after up-state cues, similarly coinciding with the first positive ERP-peak after stimulus onset, while no difference in spindle power was observable in the case of the down-state cueing. Thus, up-state cues, which were played in accordance with the endogenous rhythm of the brain, seem to recruit sleep spindles more easily than down-state cues, thereby enhancing memory consolidation in a more robust fashion. Sleep spindles are assumed to promote the re-distribution of reactivated memory representations to neocortical sites ^39^, with hippocampal reactivation signals being nested in individual spindle troughs ^4,5,40,41^. Furthermore, inducing thalamic sleep spindles, when phase-locked to SO up-states enhances the oscillatory coupling between SOs, spindles and hippocampal ripples and furthermore memory consolidation ^42^. However, memory reactivation processes in rodents seem to slightly precede the appearance of sleep spindles ^43^, while cortical sites are presumably even shut off from hippocampal inputs during the presence of sleep spindles ^44^. This might suggest that sleep spindles themselves rather enable locally undisturbed cortical reprocessing of reactivated memories ^45^. Still, there is little doubt that sleep spindles are tightly linked to slow oscillatory activity and consequently memory processing during sleep.

Bolstering this assumption, Ngo and colleagues ^46,47^ have previously demonstrated that entraining SOs through closed loop auditory stimulation, enhances phase-locked spindle activity and importantly memory recall after sleep. Interestingly, the associated increases in sleep spindle amplitude were positively correlated with later memory performance. These studies were able to elegantly demonstrate that elevating activity in the SO and phase-locked sleep spindle range by stimulation of SO up-states leads to a general improvement in memory performance. However, whether these effects resulted from specifically enhancing reactivation processes through up-state stimulation remained unknown. In our study, we are able to test this relation more directly by targeting the memory content specifically into the proposed functional up-state of the SO and compare this to TMR in the SO down-state. As pointed out above we also found an increase in spindle power for up-state cues when contrasting later remembered and non-remembered words, providing a more direct link between SO phases, spindles and memory performance. The SO phase-specificity of sleep spindles has been shown consistently in the existing literature (e.g. ^5,48^), as well as their conduciveness for the stabilization of memories ^49,50^. In a recent study, Weigenand and colleagues ^51^ induced slow oscillations by presenting repeated tones in an open-loop stimulation paradigm. Here, an initial tone was played into a random SO phase, evoking a SO or K-complex with high probability and re-setting the SO to a known phase. Subsequent tones where then played into SO up-states. Interestingly, this approach led to decreases in spindle power and no beneficial effect on memory performance compared to a control condition. Similarly, in our experiment, when presenting word-cues during SO down-states we forced the signal into a slow oscillatory rhythm, while neither a memory specific spindle response nor a stable memory enhancing effect became apparent. This might further indicate that inducing a SO out of phase has a reduced probability of triggering memory consolidation relevant processes.

In sum, our results suggest that slow-oscillatory up-states present an optimal time window for benefitting memory by TMR. Still, it must be noted that the impact of up-state associated TMR did not exceed the usually described ~10 percent benefit of memory cueing in previous ‘random-phase’ TMR studies ^17,19^. The equivalence concerning the obtained effects might result from the fact that those earlier studies featured a high stimulus repetition rate (~10 repetitions per memory cue compared to ca. 5 repetitions in the current study). In addition, presenting memory cues at suboptimal phase appears to slightly support the consolidation of memories according to our findings, especially when followed by subsequent REM sleep. As random-phase TMR allows for higher stimulus repetitions and is possibly easier to implement, it might even be the method of choice for enhancing memories during nighttime sleep in real-life.

## Materials & Methods

### Subjects

A total of 22 healthy, right-handed subjects (18 female, mean age = 20.85 ± 0.28) with German mother tongue and without Dutch language skills participated in the study. 6 subjects had to be excluded from the study due to technical reasons (n = 5) or because the subjects were too sensitive to the auditory cues and could not sleep (n = 1).

None of the participants was taking any medication at the time of the experiment and none had a history of any neurological or psychiatric disorders. All subjects reported a normal sleep-wake cycle and none had been on a night shift for at least 8 weeks before the experiment. On experimental days, subjects were instructed to get up at 7:00h and were not allowed to consume caffeine or alcohol or to nap during daytime. All participants spent an adaptation night in the sleep laboratory prior to the experiment. The ethics committee of the Canton of Fribourg approved the study, and all subjects gave written informed consent prior to participating. After completing the whole experiment, participants received 120 Swiss Francs or course credit for participating in the study.

### Design and Procedure

Participants entered the laboratory at 21:00h. The session started with the application of the electrodes for standard polysomnography, including electroencephalographic (EEG; 32 channels, Brain Products GmbH), electromyographic (EMG), and electrocardiographic (ECG) recordings.

The encoding phase started at ∼22:00 h with the learning of Dutch-German word pairs (for a detailed description see ‘Vocabulary Learning Task’ section). After completing the learning task participants went to bed at 23.00 h and were allowed to sleep for 3 h. During the 3-h retention interval, a selection of the prior learned Dutch words was presented again during sleep stages N2 and SWS. At ∼2.00 h, subjects were awakened from sleep stage 1 or 2 and recall of the vocabulary was tested again (see Fig. 2a).

### Vocabulary-Learning Task

The vocabulary-learning task consisted of 120 Dutch words and their German translations. There were three learning rounds. In each, Dutch words were presented aurally (duration range 300–500 ms) via loudspeakers (70 dB sound pressure level). In the first learning round, each Dutch word was followed by a fixation cross (500 ms) and subsequently by a visual presentation of its German translation (2000 ms). The inter-trial interval between consecutive word pairs was 2000 – 2200 ms. Subjects were instructed to memorize as many word pairs as possible. In a second round, the Dutch words were presented again followed by a question mark (ranging up to 7 seconds in duration). The participants were instructed to vocalize the correct German translation if possible or to say, “next” (German translation: “weiter”). Afterward, the correct German translation was shown again for 2000 ms, irrespective of the correctness of the given answer. In the third round, the cued recall procedure was repeated without any feedback of the correct German translation. Recall performance of the third round (without feedback) was taken as pre-retention learning performance. Here, participants recalled on average 60.5 ± 11.29 words (range 43 to 82 words) of the 120 words correctly, indicating an ideal medium task difficult (50.42% words remembered).

### Reactivation of Vocabulary

Of the 120 words learned before sleep, 2/3 of the remembered and 2/3 of the non-remembered words, totaling 80 words, were randomly selected for cueing during sleep. The remaining 40 words were not replayed during sleep (mean remembered uncued: 20.56 ± 0.95). From all words selected for cueing, half of the remembered and half of the non-remembered words were randomly selected for up-state cueing and the other half for down-state cueing (mean remembered up-state: 19.88 ± 0.90; mean remembered down state: 20.00 ± 0.95 down-state). Cueing of the Dutch words started after the participants entered stable N2 sleep and was paused as soon as arousals were detected. During the 3 hour retention phase words were presented aurally via loudspeakers (55 dB sound pressure level) either during the up-state of a SO (up-state cueing) or during the down state of a SO (down-state cueing) for a total of 90 minutes.

### Online Detection Algorithm

The open-source FieldTrip toolbox ^52^ was used to accomplish the online detection of slow oscillations and auditory replay. Slow oscillations were detected on the basis of EEG recordings from electrode site Fz, because SOs usually originate in the prefrontal cortex ^53^. Post-hoc analysis of the phase distribution at cue onset shows an even phase distribution for all channels (See Supplementary Fig. 3). The signal was referenced to the average potential from linked mastoid electrodes and filtered between 0.2 and 4 Hz. A custom fieldtrip script running under Matlab enabled to respond to the incoming EEG data from electrode Fz in real time. To detect up-and down-states, respectively, each time the EEG signal crossed an adaptive threshold the auditory replay was triggered. The algorithm was implemented as a finite-state machine (see Supplementary Fig. 1): In the first state the algorithm tries to detect a potential slow wave by waiting for the EEG signal to go below −75 µV ((1) in Fig. 1a). If the auditory stimulus is to be played in the upstate, the presentation is triggered as soon as the EEG signal goes back above −75 µV (see position (2) in Fig. 1a). On the other hand, if the stimulus is to be released during a down-state, the following state will wait for the signal to pass into the positive domain (above 10 µV) ((3) in Fig. 1a) and then again go below the release threshold of 10 µV ((4) in Fig. 1a). At this point the auditory stimulus is played. As the duration of all words was ~400 ms, they could fit within their respective target state. After triggering the replay, an 8 second recovery period is entered before the algorithm returns to its initial state.

### Recall of Vocabulary after the Retention Interval

During the recall phase, the 120 Dutch words were presented again aurally in a randomized order. The participants were asked to vocalize the correct German translation if possible. As index of memory recall of German translations across the retention interval, we calculated the relative difference between the number of correctly recalled words before and after the retention interval, with the pre-retention memory performance set to 100%.

### Sleep EEG

Sleep was recorded by standard polysomnography including EEG, electromyographic (EMG) and electrocardiographic (ECG) recordings. EEG was recorded using a 32-channel system (EasyCap, Brain Products GmbH). Impedances were kept below 10 kOhm. Voltage was sampled at 500 Hz and initially referenced to the vertex electrode (Cz). In addition to the online identification of sleep stages, polysomnographic recordings were scored offline by 3 independent raters according to standard criteria^54^.

### Preprocessing

EEG preprocessing was performed using Brain Vision Analyzer software (version 2.1; Brain Products, Gilching, Germany). Data were re-referenced to averaged mastoids and low passed filtered with a cutoff frequency of 30 Hz. The data was segmented into 6 second segments, beginning 3000 ms before stimulus onset. Trials including artifacts (e.g. movement artifacts) were manually removed after visual inspection. Afterwards, epochs were categorized into up-and down-state stimuli, depending on whether they were presented during up-or down-states, respectively. Furthermore, all stimuli were differentiated on a behavioral level into ‘Remember’ and ‘Non-Remember’ words. Remembered words refer to those words that were remembered at recall after sleep, while non-Remember words were not. Additionally ‘Up-All’and ‘Down-All’ will be used to denote all up-state and all down-state cues irrespective of the behavioral outcome. All further analyses were done using Matlab (The Math Works Inc., Natick, MA, USA) and the FieldTrip toolbox ^52^.

### Event-Related Potentials

ERPs were analyzed using FieldTrip. Trials were averaged and baseline corrected for each stimulus category within each subject. For baseline correction the segment from −3 to −2 seconds was used, as the pre-stimulus time window closer to the cue-onset systematically differs between up-and down-state. Subsequently the ERPs were averaged across all subjects for each stimulus category.

### Slow Wave Phase Analysis

The preprocessed data was low-pass filtered at 1.5 Hz and a Hilbert transform was applied. The angle information was then averaged within each behavioral category for each subject. Descriptive and inferential statistics were calculated using the Circular Statistics Toolbox ^55^.

### Time-Frequency Analysis

Time-frequency analysis was performed using FieldTrip. To obtain oscillatory power we used a continuous wavelet transformation (complex Morlet waveform, 5 cycles) for frequencies ranging from 0.5 to 20 Hz, in steps of 0.5 Hz and 10 ms. The frequency data was normalized using data from −3 to −2 seconds pre-stimulus as baseline for each stimulus category. Then the trials were averaged for each subject.

### Statistical Analysis

We analyzed the behavioral data using MANOVAS. We used post hoc paired t-tests corrected for two-sided testing. Pearson’s linear correlation coefficient was computed. A threshold of *P* = 0.05 was used to set statistical significance. For the time-frequency analysis we tested the difference between the remembered and non-remembered words with a cluster based permutation test with dependent samples and a cluster level alpha of 0.05. Monte Carlo p-values were computed on 1000 random data partitions. The critical alpha-level was set to 0.05. We first tested for significant clusters broadly from 0 to 2 seconds after stimulus onset, across all channels and across all frequencies (0.5 to 20 Hz), correcting for two-sided testing. In a next step, we specifically tested the frequency bands of interest (i.e. averaged theta power: 5 to 8 Hz; averaged spindle power: 11 to 15 Hz) for positive clusters, as both frequency bands have been shown to be related to cueing success in previous studies ^12,17–19,36,37^. Testing was done independently for up-as well as down-state trials. We also tested Up-All versus Down-All to obtain the general oscillatory differences between the up-state and down-state TMR.

## Author Contributions

T.S., M.G. and B.R. designed the experiment, E.vP. and T.S. carried out the experiments, M.G.,T.S. and E.vP. analyzed the data and M.G., T.S. and B.R. wrote the paper.

## Acknowledgements

B.R. is supported by the Swiss National Science Foundation (SNSNF) (100014_162388) and the Clinical Research Priority Program (CRPP) “Sleep and Health” from the University of Zurich.

T.S. is supported by a grant of the Swiss National Science Foundation (SNSF) (P2ZHP1_164994).

